# RhizoBindingSites v2.0 is a bioinformatic database of DNA motifs potentially involved in transcriptional regulation deduced from sites of the itself genome

**DOI:** 10.1101/2024.02.22.581308

**Authors:** Hermenegildo Taboada-Castro, Alfredo José Hernández-Álvarez, Jaime A. Castro-Mondragón, Sergio M. Encarnación Guevara

## Abstract

RhizoBindingSites is a *de novo* depurified database of conserved DNA motifs potentially involved in the transcriptional regulation of the *Rhizobium*, *Sinorhizobium*, *Bradyrhizobium*, *Azorhizobium,* and *Mesorhizobium* representative symbiotic species, deduced from the upstream regulatory sequences of orthologous genes (O-matrices) from the Rhizobiales taxon. The sites collected with O-matrices per gene per genome from RhizoBindingSites were used to deduce matrices using the dyad-Regulatory Sequence Analysis Tool (RSAT) method, giving rise to novel S-matrices for the construction of the RizoBindingSites v2.0 database. A comparison of the S-matrix logos showed a greater frequency and/or re-definition of specific-position nucleotides found in the O-matrices. Moreover, S-matrices were better at detecting genes in the genome and there was a greater number of transcription factors (TFs) in the vicinity than O-matrices, corresponding to a more significant genomic coverage for S-matrices. The homology between the matrices of TFs from a genome showed inter-regulation between the clustered TFs. In addition, matrices of AraC, ArsR, GntR, and LysR ortholog TFs showed different motifs, suggesting distinct regulation. Benchmarking showed 72%, 68%, and 81% of common genes per regulon for O-matrices and approximately 14% less common genes with S-matrices of *Rhizobium etli* CFN42, *R. leguminosarum* bv. *viciae* 3841, and *Sinorhizobium meliloti* 1021. These data were deposited in RhizoBindingSites and the RhizoBindingSites v2.0 database (http://rhizobindingsites.ccg.unam.mx/).

## Introduction

The alpha-proteobacteria and two species of beta-proteobacteria are able to establish symbiosis with leguminous plants (Liu et al., 2018). When bacteria, transformed into a bacteroid form, are found within nodules in the roots of legume plants, they reduce atmospheric dinitrogen to ammonium via nitrogenase activity; this process is called symbiotic nitrogen fixation (SNF). The host plant provides the bacteroid with dicarboxylic acids to fuel the high demand for nitrogen fixation; correspondingly, the bacteroid provides this fixed nitrogen in the form of ammonium and amino acids to plant cells (Udvardi and Day, 1997). Intensified legume production based on biological nitrogen fixation instead of the predominant practice of using chemical fertilizers to cope with high food demand is a desirable and sustainable way to diminish the eutrophication of aquatic systems as well as emissions of nitrous oxide into the atmosphere (Blesh, 2019), (Erisman et al., 2011), (Ferguson et al., 2019).

A better understanding of the role of bacteroid transcription regulators (TFs) in SNF processes is fundamental for designing a better genetic regulatory circuitry to enhance the ability of symbiotic bacteria to reduce dinitrogen (Marx et al., 2016). This requires great effort because *Rhizobium* is a free-living and symbiotic bacterium. Regulatory circuits with intricate wiring have evolved to adapt to different environments (Ibarra-arellano et al., 2016) (Taboada-Castro et al., 2022), with symbiotic types, which are carried out with different leguminous plants, showing different types of nodules (determined and undetermined) (Liu et al., 2018).

High-throughput approaches, such as transcriptomics, proteomics, and metabolomics, have been applied to study the symbiosis of symbiotic species (Lardi and Pessi, 2018). Recently, metabolic maps of carbon-, nitrogen-, and phosphorus-integrating plant cells of legume nodules and bacteroids have been reported (Liu et al., 2018). Moreover, a proteomic atlas of the host plant *Medicago truncatula* and its symbiont *S. meliloti* was constructed (Marx et al., 2016). In addition, methods for annotating TF-binding sites and motif databases have been constructed, such as RegPrecise 3.0 (https://regprecise.lbl.gov/) (Novichkov et al., 2013) and RhizoBindingSites, (see below) (http://rhizobindingsites.ccg.unam.mx/) (Taboada-Castro et al., 2020). The next step would be to propose genetic circuits to build a transcriptional regulatory network for SNF. A bioinformatics method to construct *in silico* transcriptional regulatory networks, comparing the free-life and symbiosis in the maximal nitrogen fixation stage from *R. etli* CFN42 with the bean plant *Phaseolus vulgaris*, has been shown (Taboada-Castro et al., 2022). In addition, a computational reconstruction of the transcriptional regulation of nitrogen fixation and signaling by oxygen in alphaproteobacteria was reported (Tsoy et al., 2016). Although the main TFs has been identified experimentally, the lack of information on the functions of TFs related to carbon, phosphorus, and minerals, among others, makes it difficult to integrate them into a network. It is necessary to promote experimental designs that describe the role of a complete set of genes expressed in response to stimuli. The main goal of this work is to Provide computational information on gene regulation for experimentalists for a more precise design of experiments.

We recently released the RhizoBindingSites database (http://rhizobindingsites.ccg.unam.mx/). These data were depurified (see Materials and methods); the RhizoBindingSites database was obtained with the phylogenetic footprinting algorithm (Regulatory Sequences Analysis Tools, RSAT) footprint-discovery (Janky and van Helden, 2008), which aligns the promoters of orthologous genes for each genome in the order Rhizobiales to deduce a position-specific scoring matrix (PSSM), which represents the motif. These motifs are composed of spaced sites or dyads that are conserved among species and are potentially recognized by TFs. The dyads are two to three conserved nucleotide sequences spaced by a non-conserved sequence (separator) located in the regulatory region of the gen. The matrices, hereinafter referred to as “orthologue-derived-matrices” (O-matrices), were used to scan gene promoters in a genome to give rise to hypothetical regulons (h-regulons), which are defined as a group of genes sharing a motif of a TF, but, conventionally, non-TF genes also potentially have a motif in common. The output table (RhizoBindingSites, Motif Information section) contains an h-regulon per gene per genome with additional information (Taboada-Castro et al., 2020).

In this report, all the “sites” corresponding to the motifs of an h-regulon, in the Motif Information section of RhizoBindingSites in the “sites” column (http://rhizobindingsites.ccg.unam.mx/), were used to newly predict matrices hereinafter referred to as “single-genome-matrices” (S-matrices). Remarkably, the RhizoBindingSites O-matrices originated from diverse orthologous genes from different species. In contrast, the S matrices were deduced from the sites of the respective genomes. The S-matrices (as was shown for the O-matrices) (Taboada-Castro et al., 2020) were used to scan the upstream −400 to −1 regulatory regions of all genes in their respective genomes, such as *R. etli* CFN 42, *R. etli* bv. *mimosae* Mim1, *Bradyrhizobium diazoefficiens* USDA 110, *Sinorhizobium fredii* NGR234, *Sinorhizobium meliloti* 1021, *R. l.* bv. *viciae* 3841, *Bradyrhizobium* sp. BTAi1, *Azorhizobium caulinodans* ORS 571, and *Mesorhizobium japonicum* MAFF303099, giving rise to RhizoBindingSites v2.0. We noticed that S-matrices contained more genes than O-matrices in their respective genomes. A comparison of logos showed that the frequency of nucleotide position specificity was better for S-than for O-matrices, but for other motifs, re-deduction of matrices yielded different nucleotide compositions. A vicinity analysis of the genes per genome detected with S-matrices included more TFs than those detected with O-matrices. A matrix-clustering analysis of only matrices of TFs per genome for O-and S-matrices showed clusters of TFs with a minimum of two different genes, suggesting a functional relationship and different regulation for TFs of the same family.

## Material and Methods

All bioinformatics methods used to construct the RhizoBindingSites database were performed using RSAT (http://embnet.ccg.unam.mx/rsat/) in a Linux environment (Taboada-Castro et al., 2020).

### Deduction of O-matrices in the RhizoBindingSites database

PSSMs or O-motifs were deduced using the footprint discovery RSAT algorithm (Nguyen et al., 2018). Briefly, this program receives, as input, the name of an organism, one or more gene names, and the name of a taxon (command in Appendix A). The program searched for orthologous genes in the given taxon for each gene of the desired organism with the best bidirectional hit and an E-value < 1.0e-5. For each selected ortholog, the program obtained the upstream sequences (−400 to −1) with respect to the translation start site. These sequences were masked in redundant fragments and a motif discovery algorithm called dyad-analysis-RSAT (van Helden et al., 2000) was applied to these sequences. This program looks for over-represented variable-spaced motifs, which is the case for most bacterial TFs that bind to DNA as dimers. In addition, the motif is represented by a short oligonucleotide. The detected dyads or oligos were assembled into PSSMs. For this analysis, we used a background model, taking all upstream sequences of all organisms belonging to the order Rhizobiales (Brohée et al., 2011) (Taboada-Castro et al., 2020).

### Deduction of S-matrices in the RhizoBindingSites v2.0 database

The sites located in the Motif Information section of the RhizoBindingSites database, created by the genomic matrix-scan with the filtered O-matrices from the RhizoBindingSites database was extracted, these nucleotide sequences representing the motifs were used as input to the program “create_matrices_from_matches_2018. pl” (Appendix A). This program generates two files: one with sites per predicted regulon and one with purged sites with the .fas extension in the new directory “Binding_sites_files.” This directory is used as input to the program “motifs_discovery_from_matches_sequences_2018. pl” (appendix B). This program deduces the matrices using a PSSM with the dyad-analysis method described previously (Taboada-Castro et al., 2020), creating a new directory called “motif_discovery” that contains one sub-directory per h-regulon generated for each gene (the directory name is the same as the gene that gave rise to the regulon), which contains three files: dyads, the pattern assembly of the dyad, and one to five matrices in the transfac format with the .tf extension.

### Filtering of matrices

The S-matrices were filtered by selecting those able to find motifs in their own promoter gene through the matrix-scan RSAT program at a p-value ≤ 1e-4 as was shown for O-matrices (Taboada-Castro et al., 2020) (Defrance et al., 2008) (Thomas-Chollier et al., 2008).

### Genomic matrix-scan for selecting targets with S-matrices

The S-matrices were used to scan the-400 to the −1 promoter region of all genes of their respective genome through the matrix-scan RSAT program with a p-value ≤ 1e-4 as was described (Taboada-Castro et al., 2020). These output data contained the h-regulons predicted with the S-matrices that were conventionally fractionated to stress the importance of having data with different levels of stringency: low (p-value: 1.0e-4 to 9.9e-4), medium (p-value: 1.0e-5 to 9.9e-5), and high (p-value: 1.0e-6 to lowest data value). Stringency is referred in the scanning process to the homology between the nucleotide sequence of the S matrix and the sequence in the upstream regulatory region of genes in both the forward and reverse strands of DNA. These data define a hypothetical regulon (h-regulon) per gene, which is a group of genes in a genome detected during the scanning process with S-matrices of the aforementioned gene (Taboada-Castro et al., 2020). Since matrices detect motifs in both DNA strands through matrix-scan analysis, additional *de novo* depuring step data were applied by selecting only targets in the codificant string of the target gene for O- (RhizoBindingSites) and S-matrices (RhizoBindingSites v2.0).

### Motif information and gene information sections

Genomic matrix-scan analysis with the filtered S-matrices generated the motif information data similar to the Motif Information section of the RhizoBindingSites database. Gene Information sections were as in the RhizoBindingSites database (Taboada-Castro et al., 2020).

### Vicinity of genes from an h-regulon

For bacterial genomes, proximal genes are often functionally related; this may occur because they may be in the same operon and regulated by the same TF (Bowers et al., 2004) (Pannier et al., 2017) (Strong et al., 2003), but may also be genes with their promoters. Vicinity was defined as a group of genes at a distance of less than or equal to three genes in the genome (Taboada-Castro et al., 2020). The vicinity was searched for each gene in the genome and genes grouped in the COGK category (Tatusov et al., 2003) (Appendix C) related to transcriptional regulation, that is, TFs, response regulators, two-component response regulators, sigma factors, and anti-sigma factors. At low, medium, and high levels of p-value stringency, as shown in the Gene Information section of the RhizoBindingSites database (Taboada-Castro et al., 2020), a comparison of the percentage of TFs with neighbors of h-regulons from the data of the O- and S-matrices at the three p-values is shown (Supplementary Table 1C).

### Matrix-clustering

S- and O-matrices from only the COGK genes (Tatusov et al., 2003) per genome were selected for matrix-clustering RSAT analysis (Appendix D) (Castro-Mondragon et al., 2017). This program constructs clusters by grouping S- or O-matrices based on similarity. There is an output file “clusters_motif_names.tab” containing the clusters in a list (i.e., cluster_3 RHE_RS06555_m1, RHE_RS06555_m2, RHE_RS03090_m3, RHE_RS03090_m1 and RHE_RS03090_m2”).

Genes RHE_RS06555 and RHE_RS03090 appeared repeatedly in this cluster. To avoid this redundancy, unique genes per cluster were counted using the clusters_motif_names.tab file, and clusters with at least two different genes were extracted. Subsequently, NCBI information for each gene was added (Supplementary Table 2).

### Comparison of regulons from the Regprecise database, RhizoBindingSites, and RhizoBindingSites V2.0

We referenced the Regprecise data because it included data from 56 regulons of the studied species in the RhizoBindingSites and RhizoBindingSites v2.0 databases. This was constructed using the experimental data from CoryRegNet 4.0, RegTransBase, and Regulon database (V6.0) (Novichkov et al., 2013). Briefly, data were obtained by searching for TFs and their target gene orthologs in a group of representative phylogenetically related species. Ortholog TFs with their target genes are called regulogs, and the search for regulogs was extended to the rest of the species of a taxon. A position weight matrix was used to deduce the motifs for each regulator using their methodology. The authors considered this data propagation to be accurate and conservative, indicating that it was not an attempt at automatic prediction (Novichkov et al., 2013).

In contrast, our data were *ab initio* deduced from orthologous genes without a reference. First, the equivalent locus tags of the regulons of the extended section of Regprecise from *Rhizobium etli* CFN42, *R. leguminosarum* bv. *viciae* 3841, and *Sinorhizobium meliloti* 1021 were searched for compatibility with our locus tags (Supplementary Table 7A–C). Then, we used the application from our databases, “Prediction of regulatory networks,” by pasting the TF to the left box and the potential targets to the right box with the option “auto”; each of the 56 regulons was reported in the propagated section of Regprecise 3.0 (https://regprecise.lbl.gov/), for both O- and S-matrices (data not shown). Since, in our data, the operon arrangement of the genes was not considered and most of the Regprecise regulons were operons, a paired list was constructed between the Regprecise operons and the genes found with the O- and S-matrices in our databases. Only the genes of regulons with genes of O- and S-matrices were considered common genes (Supplementary Table 7A–C).

### Useŕs Guide

RhizoBindingSites and RhizoBindingSites v2.0 are intuitive databases, additionally a Useŕs guide is included, a Matrix-clustering window and Synonyms converter application were included. For matrix-clustering data, given that a gene may have from one to five matrices, these matrices frequently clusters, giving rise to clusters from the same locus-tag, which is low informative, to solve this, a data with clusters, containing more than two different genes may be consulted as a guide search (Supplementary Table_2_Matrix-clustering_Analysis_of_O_and_S-Matrices). Additionally, for the application “Prediction of regulatory Networks” the user needs to use the locus tags proper of the application, to correct this, the synonyms converter was implemented (http://rhizobindingsites.ccg.unam.mx/).

## Results and discussion

### Statistics of RhizoBindingSites (O-matrices) and RhizoBindingSites v2.0 (S-matrices)

We showed the relevance of considering the stringency level of the inferred data on transcriptional regulation (see Materials and Methods). A comparative analysis of the average number of unique genes with O- and S-matrices for the three fractions of p-value showed 3871.11 and 3016.11 genes from the nine genomes, respectively (Supplementary Table 1A). On average, there were 854.7 fewer genes with deduced S-matrices than O-matrices. Correspondingly, there was, on average, 76.27% and 65.68% of TFs with O- and S-matrices, respectively, with respect to the total content of TFs in the nine genomes (Supplementary Table 1A). These data indicated that there was 11% less TF content with S-matrices than O-matrices (Supplementary Table 1A), suggesting that sites from these O-matrices had low conservation, and it was difficult to find a consensus for the re-deduction of S-matrices. Unique genes detected per genome were determined after matrix-scan analysis of the respective genomes using these matrices. These data showed, on average, taking into a count all data from low, medium and high stringency, 5455.11 and 5542.55 unique genes were detected with the O- and S-matrices per nine genomes, respectively. Furthermore, 81.63% and 82.91% of the genomic coverture concerned the average gene content of the nine genomes. There was a 1.3% greater genomic coverage for the S-matrices than O-matrices (Supplementary Table 1B), despite the fact that more genes with O-matrices were found. Given that the addition of genes due to the presence of operons was not considered in our data, the genomic coverage number could be greater than these, showing important genomic coverage with both O- and S-matrices.

### S-matrices are more accurate than O-matrices

TFs are frequently found behind gene-target neighbors (Strong et al., 2003) (Pannier et al., 2017) (Novichkov et al., 2013). We analyzed the vicinity of the h-regulons (see the Gene Information sections of RhizoBindingSites and RhizoBindingSites v2.0). Moreover, the percentage of TF content per p-value fraction per genome for neighboring genes was determined with respect to the respective TF content per genome. At the p-value fractions of 1.0e-04, 1.0e-05, and 1.0e-06 for low, medium, and high stringent data, respectively, there were seven, six, and five genomes with greater than 1.0% averages of TF content in the neighboring genes with S-matrices compared to O-matrices, respectively. Genomes *Rhizobium etli* CFN42, *Rhizobium etli* bv. *mimosae* str. Mim1, *Sinorhizobium meliloti* 1021, *Bradyrhizobium sp*. BTAi1, and *Azorhizobium caulinodans* ORS571 showed, for S-matrices, a greater TF content in the neighboring genes in the three p-value ranges. *Rhizobium leguminosarum* bv. *viciae* 3841 showed a greater TF number with neighboring genes in the 1.0e-04- and 1.0e-06-p-value ranges. In contrast, *Sinorhizobium fredii* NGR234 showed a greater TF content with neighboring genes in the 1.0e-04- and 1.0e-05-p-value ranges (Supplementary Table 1C).

Although, on average, 11% fewer TFs with S-matrices than with O-matrices, (se above) (Supplementary Table 1A), most of the genomes showed a greater TF content of the neighboring genes detected with S-matrices than with O-matrices (Supplementary Table 1C). The homology between the nucleotide sequences of the S-matrices and the upstream regulatory regions in the respective genomes was higher than that of the O-matrices, corresponding to the deduction of S-matrices with their own genomic sites. These data suggest that the S-matrices show greater accuracy than the O-matrices.

### Clusters with O- and S-matrices

The homology of the O- and S-matrices of TFs per genome was analyzed separately using a matrix clustering program (Castro-Mondragon et al., 2017) (Nguyen et al., 2018). Homology between the TF matrices was expected because of the functional relationship observed in an *Escherichia coli* K-12 transcriptional regulatory network (Martínez-Antonio and Collado-Vides, 2003). In addition, the hierarchy of TFs of minimal medium growth and symbiotic proteomes from *R. etli* CFN42 has been shown (Taboada-Castro et al., 2022). These homologies were displayed in hierarchical dendrograms using the HCLUST algorithm (Castro-Mondragon et al., 2017). Output matrix clustering data with O- and S-matrices are available at RhizoBindingSites and RhizoBindingSites v2.0, respectively (http://rhizobindingsites.ccg.unam.mx/). On the main page, there is a section called “Matrix-clustering,” which opens an archive.html that displays a new web page with some sections (dendrograms of the corresponding O- and S- matrices per genome) which are available by clicking on the “Logo Forest (dynamic browsing)” section, providing 18 directories (http://rhizobind-ingsites.ccg.unam.mx/). TFs with clustered O- and S- matrices were, on average, 61.8% and 59.3%, respectively, and the average of clusters were 206 and 184, respectively (Supplementary Table 1D– E). According to the average of TFs with O- and S- matrices (Supplementary Table 1A), clustered matrices for O- and S- matrices represented 14.47% and 6.38% fewer TFs, respectively (Supplementary Table 1D–E). These data show that there is more significant homology between the S-matrices than between the O-matrices. Data from 18 matrix-clustering analyses were extracted to avoid the redundancy of genes per cluster, clusters between other genes also contained genes with the same functions (Supplementary Table 2). These genomes contained more than 10% of genes with functional redundancy, i.e., in *R. etli* CFN42, there were fifteen identified AraC genes with deduced matrices (Supplementary Table 2) (see below Supplementary Table 3). Because an interrelationship between TFs is expected, it is important to know the functionality of O- and S- matrices by their ability to form regulons from clustered TFs.

### Clustered TFs may be functionally related

We have previously shown that genes clustered with a TF or TFs are potentially functionally organized in an hypothetical regulon (h-regulon) (Taboada-Castro et al., 2022). Some clusters were analyzed in the application from RhizoBindingSites and RhizoBindingSites v2.0 (“Prediction of Regulatory Networks”), by pasting the TF genes of a cluster from Supplementary Table 2 to both boxes; as regulators and as targets, a medium restriction level of 1.0e-05 was selected. This data showed, the TF genes of clusters with O- and S- TF-matrices are related, that is, cluster_99 and cluster_13 were for the O- and S- TF-matrices from *R. etli* CFN42, respectively. Cluster _103 and cluster_63 from *R. etli* Mim1 were for the O- and S- TF-matrices, respectively. Cluster_112 and cluster_70 from *R. leguminosarum* bv. *viciae* 3841 are for the O- and S- TF-matrices, respectively. Cluster _99 and cluster_19 were for O- and S- TF-matrices from *Sinorhizobium meliloti* 1021, respectively (Figure 1). If there is no relationship between TF-TF genes, the gene appears isolated, showing only that it recognizes a motif in its own promoter, that is for; cluster_63 from *R. etli* bv. *Mimosae* Mim1, gene RHEMIM1_RS034085, cluster_70 from *R. leguminosarum* bv. *viciae* 4841, gene RL_RS26595 (Figure 1). Bioinformatics methods include false-positive data, and a frontier challenge of bioinformatics sciences is to construct methods to diminish these data. A method was proposed to construct transcriptional regulatory networks, lowering low-stringency data with a matrix-clustering method, favoring TF gene-target relationships with conserved motifs in both the TF and gene target of the same cluster (Taboada-Castro et al., 2022). Therefore, the quality of the matrices is crucial for constructing regulons. Matrix clustering of TFs provides information for the construction of a global transcriptional regulatory network per genome.

**Figure_1.**
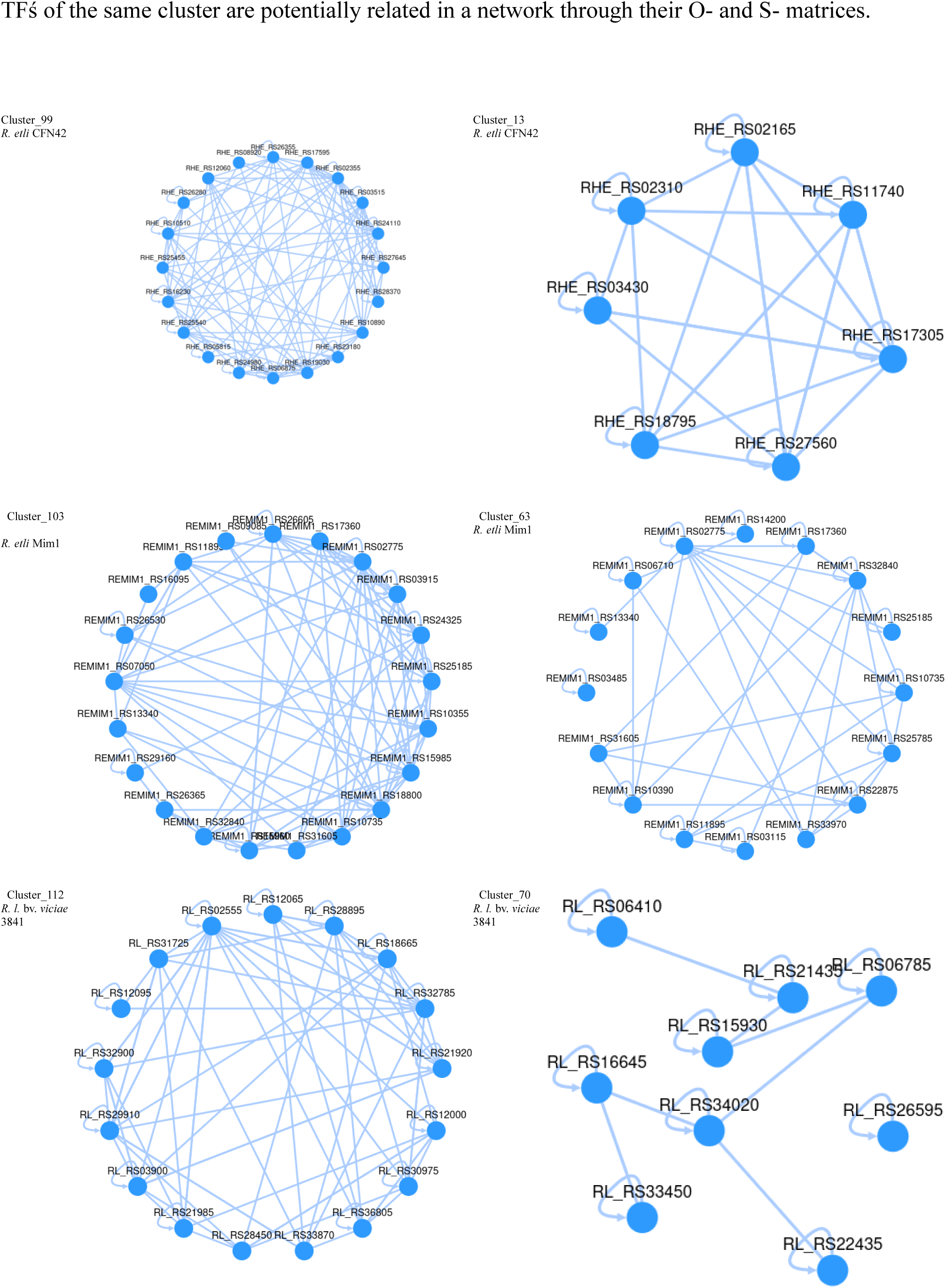
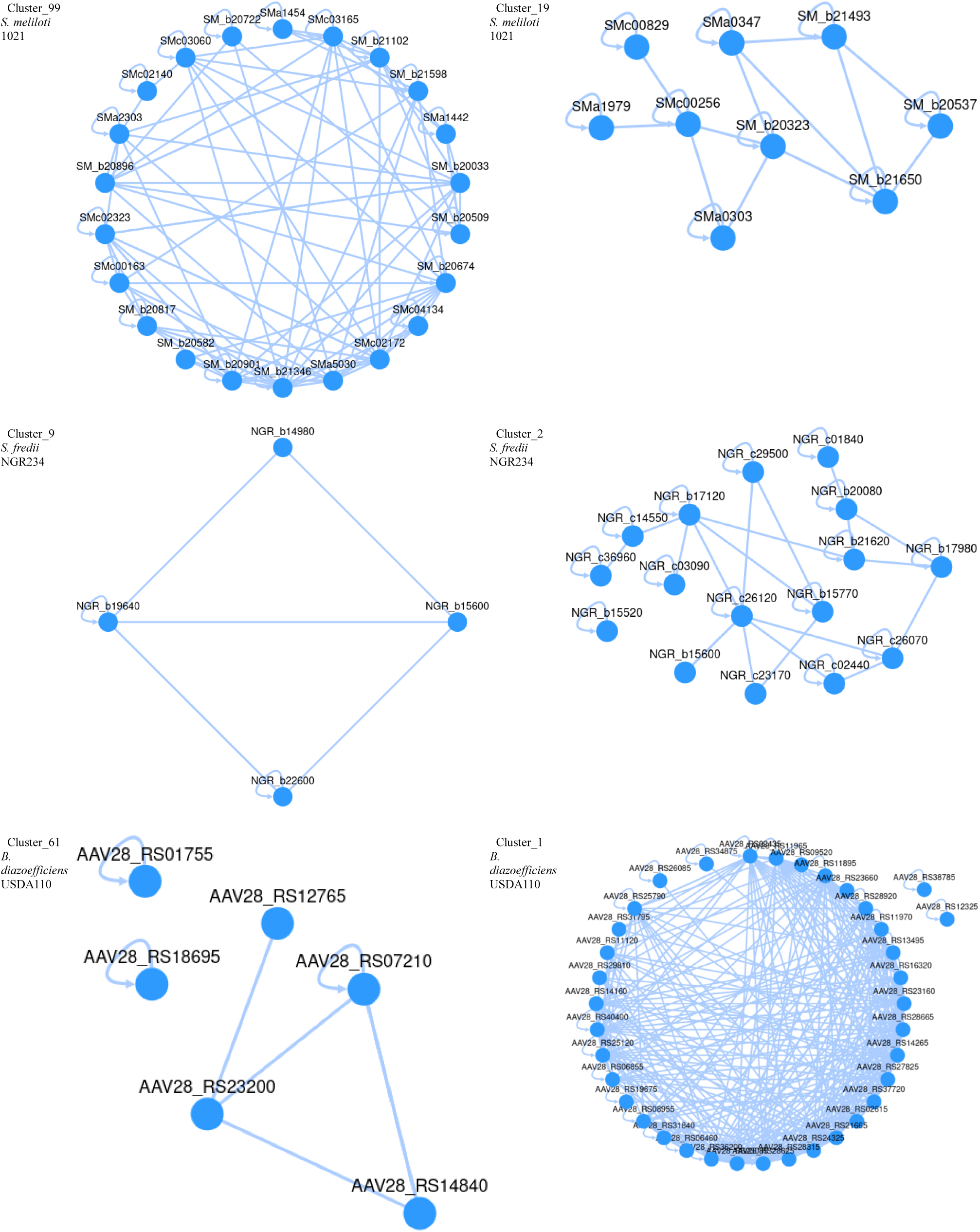
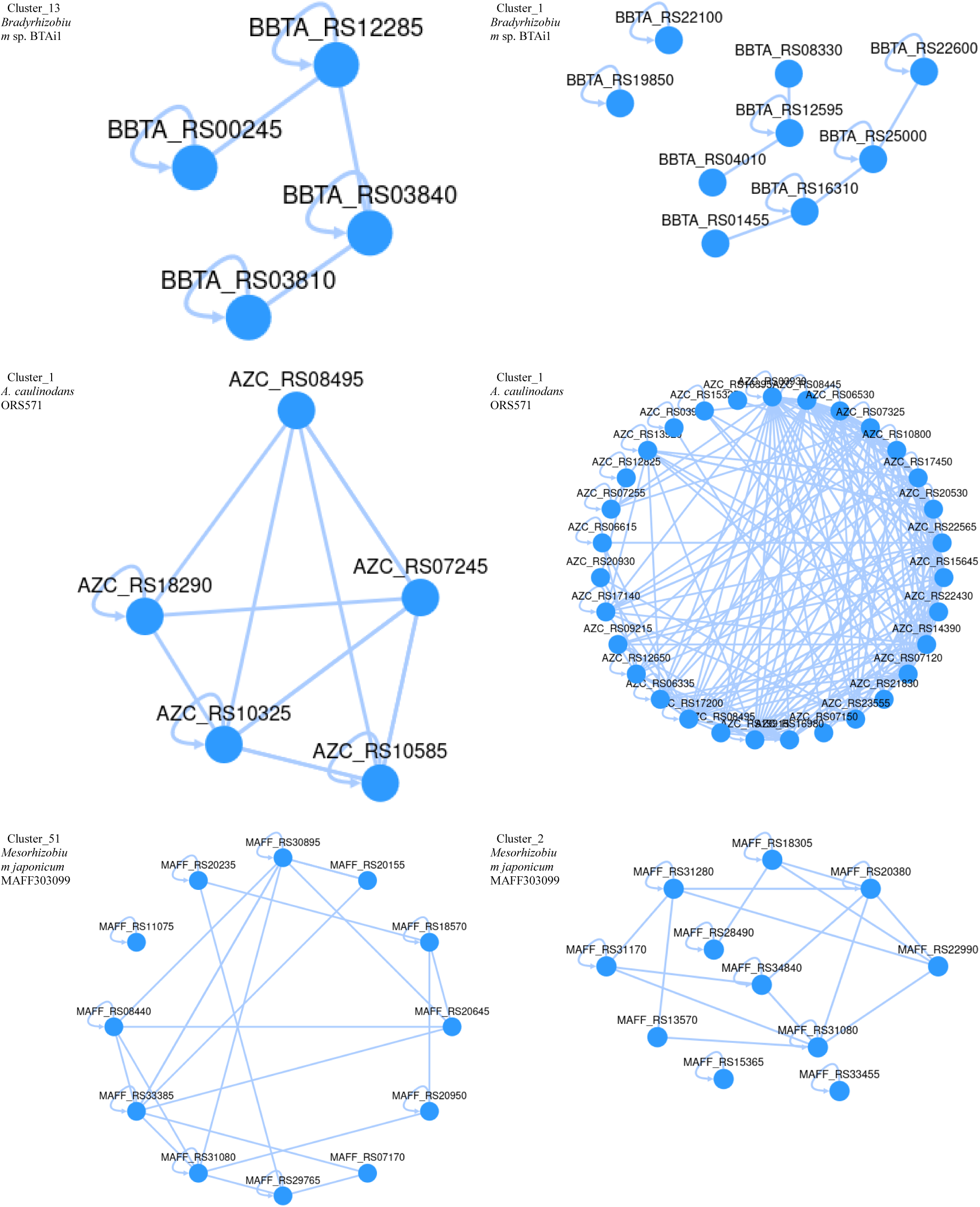
Regulons_of_TF’s. Clusters of matrices of TF-genes formed regulons in the application “Prediction of regulatory networks” for O-matrices in RhizoBindingSites and S-matrices in RhizoBindingSites v2.0. Representative clusters of TF’s from each specie were selected only for having a great number of unique genes.

### Nucleotide composition of O- and S-matrix logos

The re-definition of de matrices from sites of the genome is a novel strategy not registered in the literature. Thirty-three logos of the same gene were selected to compare the nucleotide compositions of the O- and S-matrices (Figure 2). It should be noted that cluster_83 O-matrix RHE_RS14875_m1 from *R. etli* CFN42 has a nucleotide sequence AAATTG, whereas in cluster_106, the S-matrix RHE_RS14875_m5 was replaced with AAATAT, and the sequence ACAATTT was present in both O- and S-matrices. Similarly, in cluster_51, the O-matrix RL_RS29140_m5 from *R. leguminosarum* bv. *viciae* 3841 had the sequence AAAGTGTATGCAA, as in cluster_288 S-matrix RL_RS29140_m1, but with a greater frequency in the S-matrix. In contrast, the GGACGTGCCA sequence present in RL_RS29140_m5 was replaced with TTTCG in RL_RS29140_m1.

**Figure_2.**
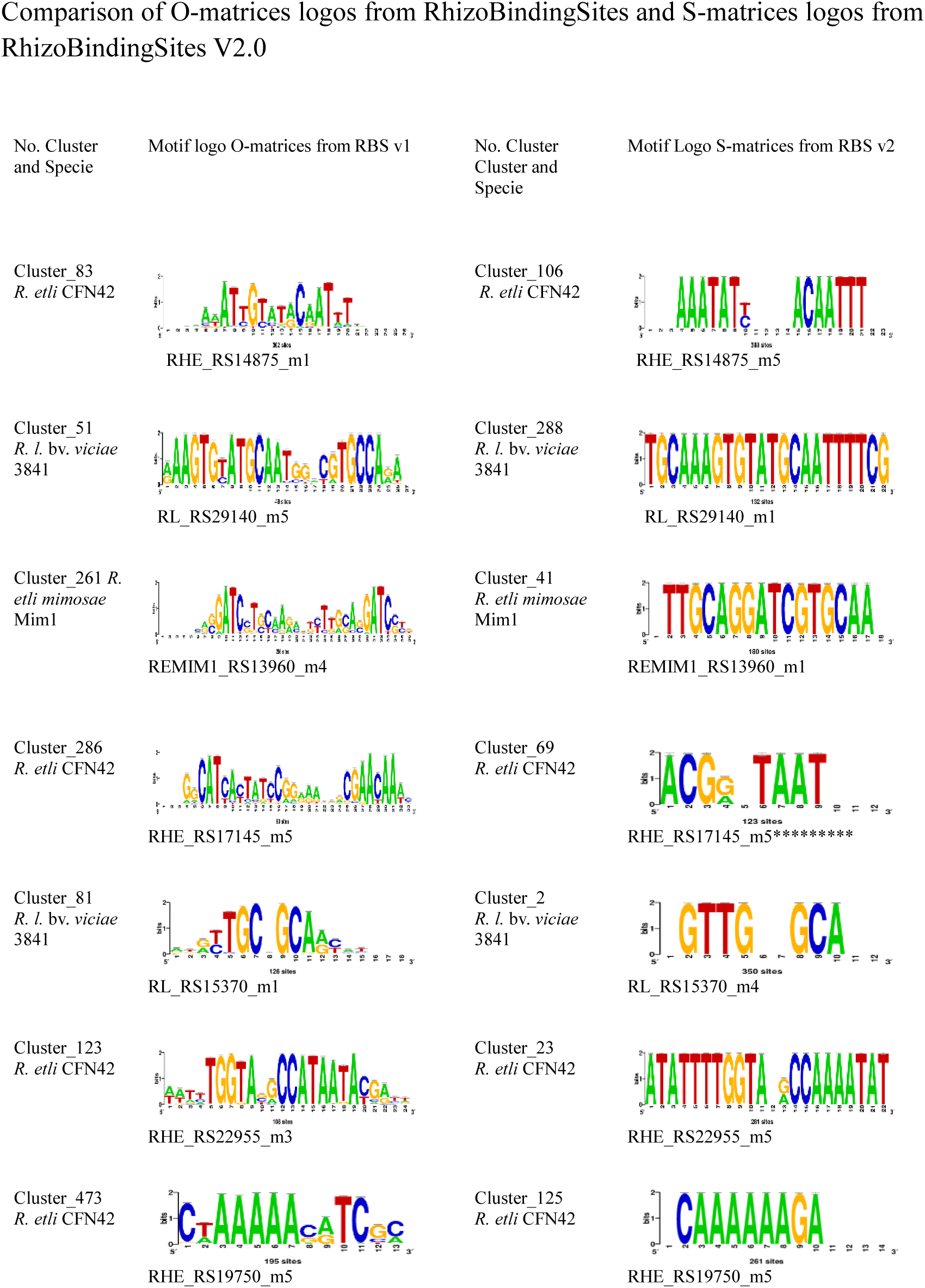
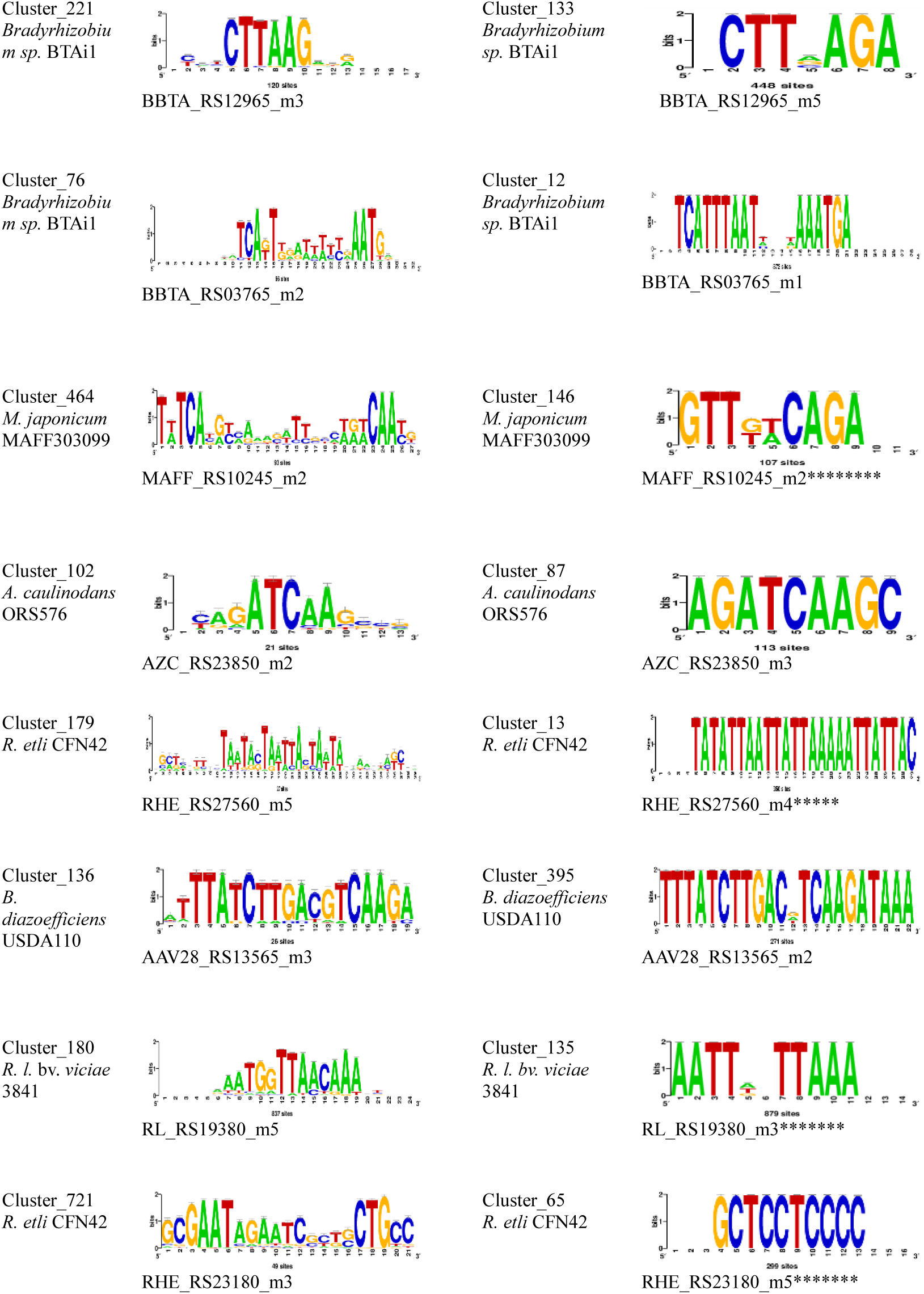
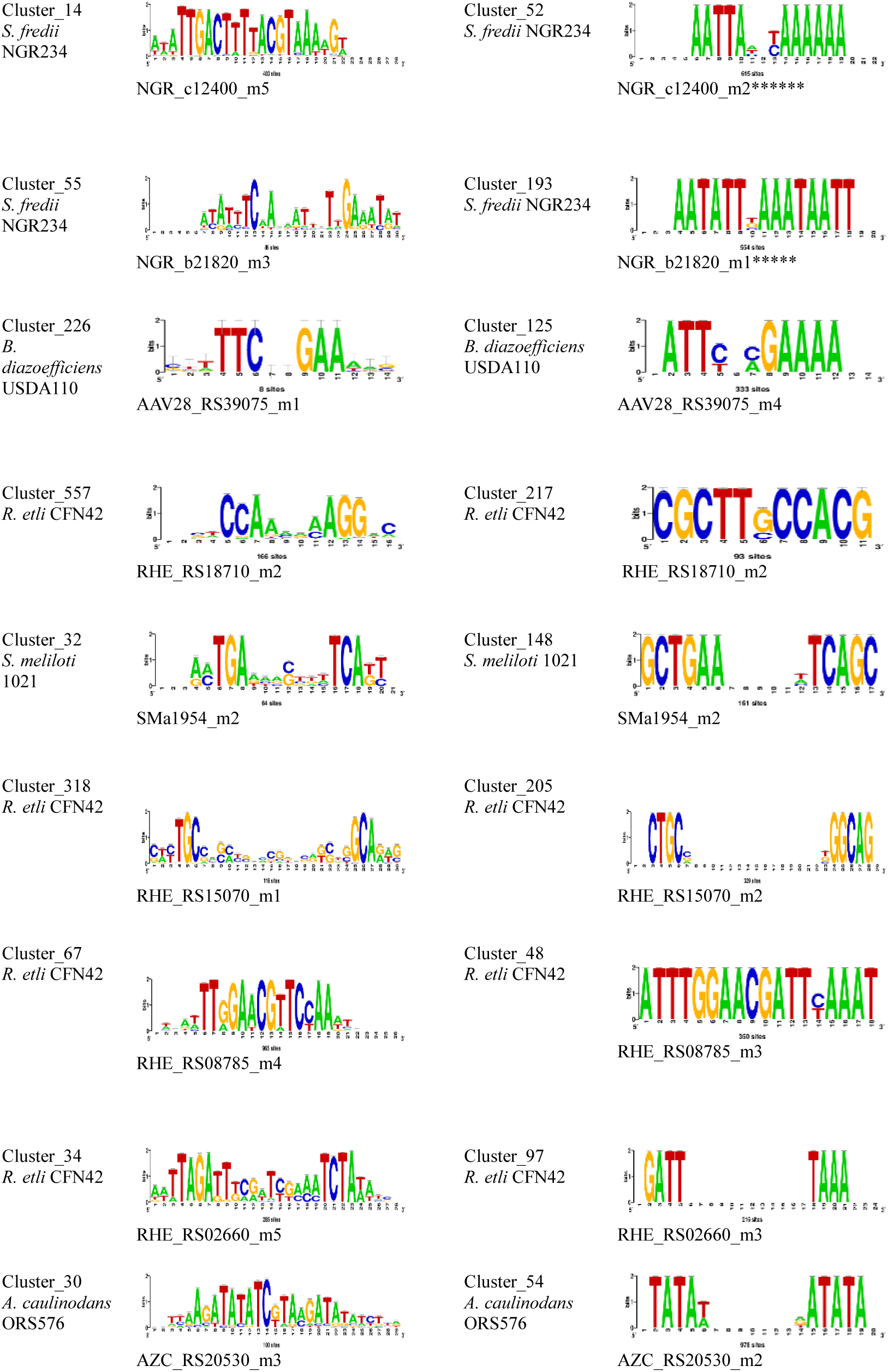
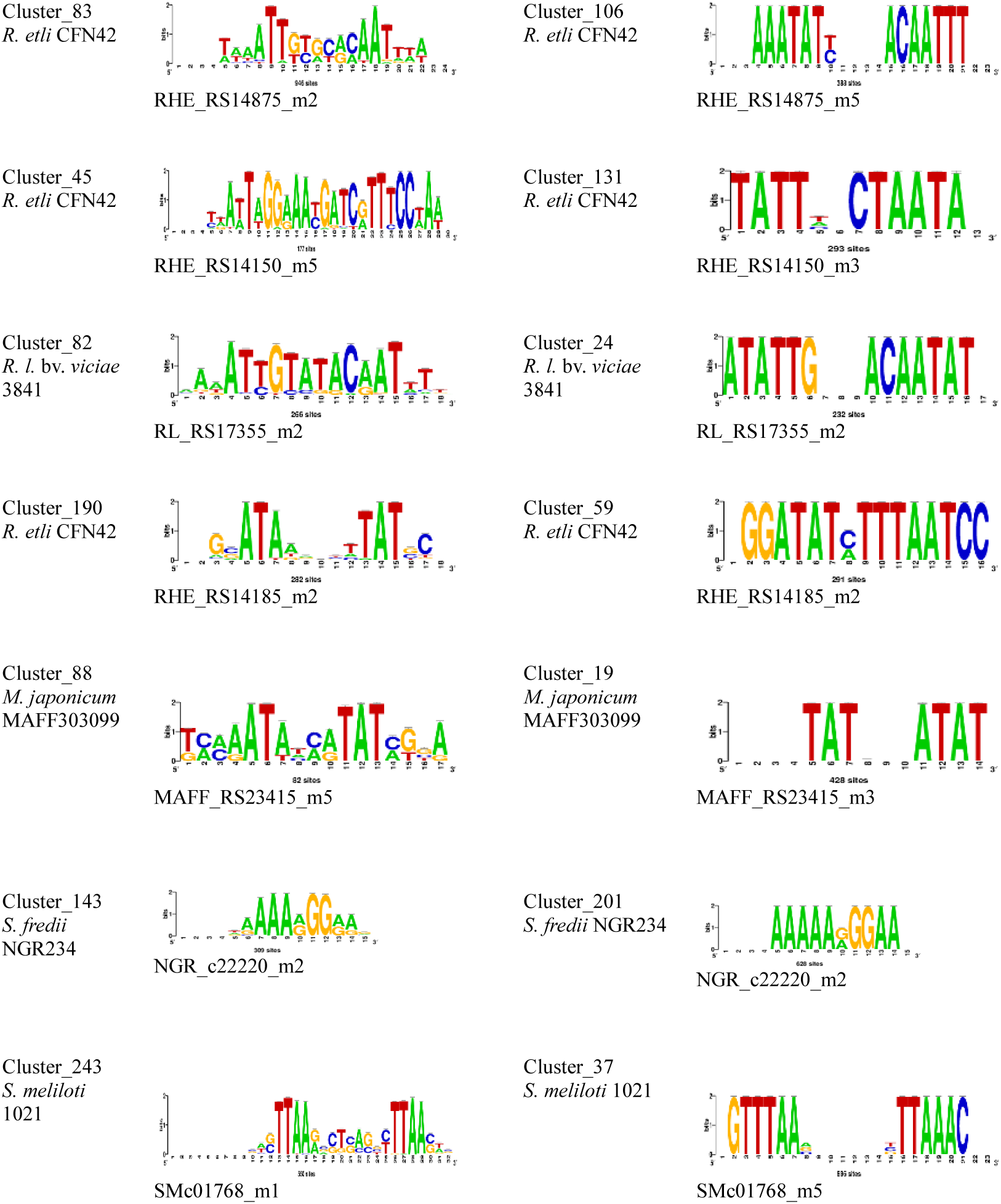
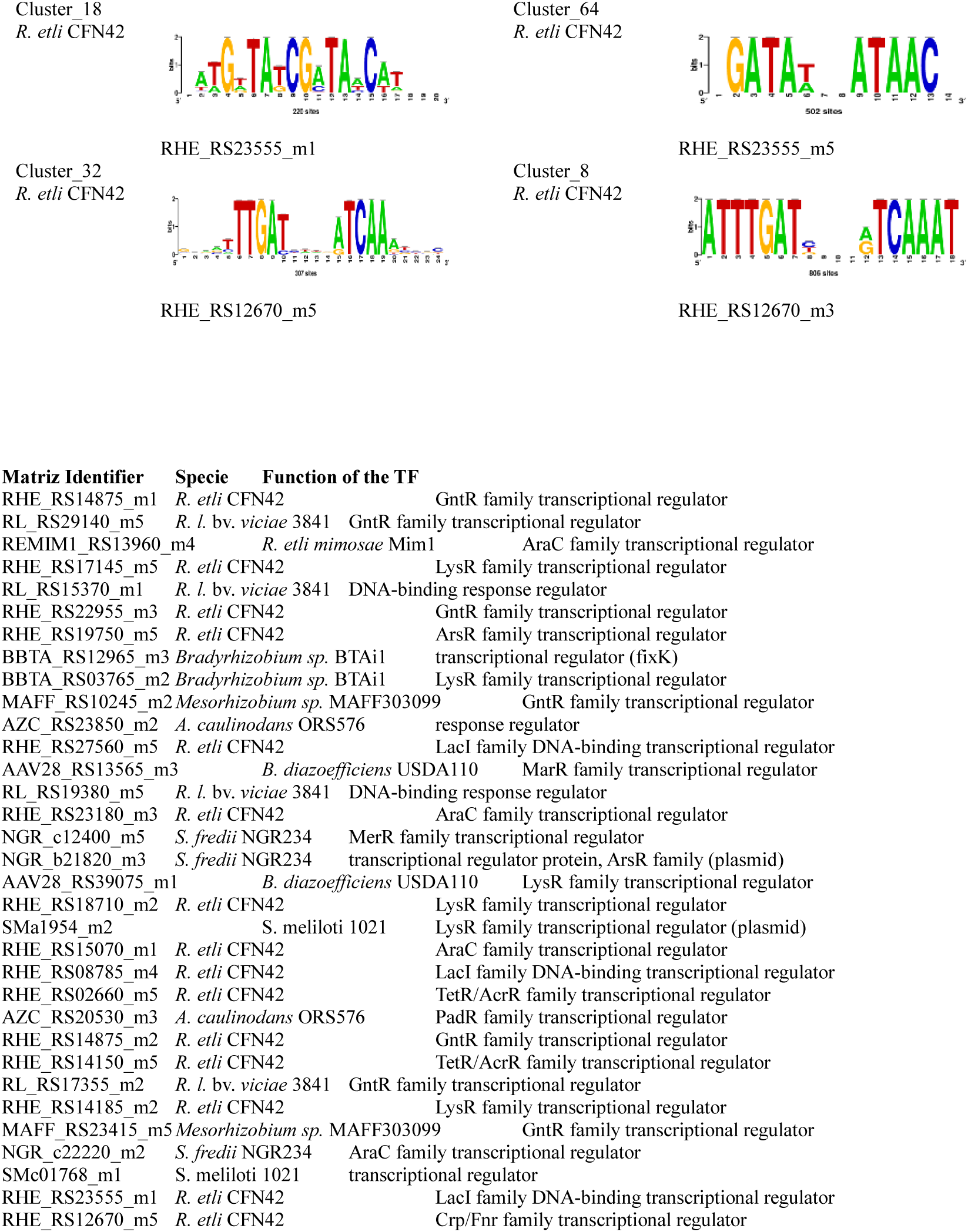
Table_of_motifs. The nucleotide composition of motifs from O- and S-matrices of the same TF was compared. The frequency of some nucleotides position specific was greater for motifs of S- than O-matrices. For other cases, the re-definition of nucleotides from motifs of S-matrices as compared with O-matrices was Observed. While, for other motifs, a completely new composition of nucleotides was observed.

Interestingly, in cluster_261, O-matrix REMIMI_RS13960_m4 from *R. etli* bv. *mimosae* Mim1, the sequence GATC-30-GATC was present, whereas in cluster_41, the S-matrix REMIMI_RS13960_m1, the oligo TTGCAGGATCGTGCAA was found, which included the GATC sequence; however, the spacing and double GATC sequence were not conserved in the sites. For cluster_286, the O-matrix RHE_RS17145_m5 from *R. etli* CFN42, and cluster_69, the S-matrix RHE_RS17145_m5 showed the sequence ACGAATAAT, but with a greater frequency in the S-matrix RHE_RS17145_m5. Moreover, the O-matrix showed the oligo GGCATCACT, which is present in the orthologs of RHE_RS17145 in the *Rhizobiales* taxon. Consequently, this oligo was present in the O-matrix but was not a consensus in the sites of the *R. etli* CFN42 genome.

From these data, we observed that some nucleotides present in the O-matrices were absent in the S-matrices. There is a re-definition of the nucleotides of the logos in the S-matrices, and the S-matrices are more suitable for their respective genomes. Around twenty-three cases with S-sites exhibited conserved nucleotide composition with O-matrices, but with a greater frequency than those in the O- matrices (Figure 2). Additionally, there was a drastic re-composition of nucleotides in the logos, i.e., cluster_286 and cluster_69 (Figure 2). Cluster _464 O-matrix *M. japonicum* MAFF303099 MAFF_RS10245_m2 and cluster_146 S-matrix *M. japonicum* MAFF303099 MAFFRS10245_m2. In addition, cluster_179 included the O-matrix *R. etli* CFN42, RHE_RS27560_m5, and cluster_13 included the S-matrix *R. etli* CFN42, RHE_RS27560_m4. Cluster_180 O-matrix *R. leguminosarum* bv. *viciae* 3841 RL_RS19380_m5 and cluster_135 S-matrix *R. l.* bv. *viciae* 3841 RL_RS19380_m3. Moreover, cluster_721 O-matrix *R. etli* CFN42 RHE_RS23180_m3 and cluster_65 S-matrix *R. etli* CFN42, RHE_RS23180_m5. In addition, cluster_14 O-matrix *S. fredii* NGR234 NGR_c12400_m5 and cluster_52 S-matrix *S. fredii* NGR234 NGRc12400_m2. This indicates, the sites obtained with the O-matrices had a lax consensus. Consequently, the re-deduction of new S-matrices from these sites was with a low consensus site, generating these differences. We notice, when AraC TFs from *R.etli* CFN42 cluster_721 O-matrix RHE_RS23180_m3 and cluster_318 O-matrix RHE_RS15070_m1 were compared (Figure 2). In addition, their corresponding cluster_65 S-matrix RHE_RS23180_m5 and cluster_205 S-matrix RHE_RS15070_m2 motifs (Figure 2), they showed different logos despite belong to the same family. This data suggested, two AraC TFs are differentially regulated. To know how much general was this observation, we analyzed more families of TFs.

### Potentially different regulation of the TFs of the AraC, ArsR, GntR, and LysR families

#### AraC family

The two AraC TFs were clustered in different groups because of the different nucleotide compositions of their motifs, suggesting distinct transcriptional regulation (see above). Matrix clustering of the O-and S-matrices of AraC TFs from *R. etli* CFN42 and *Rhizobium leguminosarum* bv. *viciae* 3841 and *S. meliloti* 1021 species were identified (Supplementary Table 3A–C). Only the AraC genes with deduced O- and S- matrices were considered. Furthermore, only clusters containing more than two different genes were considered. Note that a gene may appear more than once in the same cluster because each gene may have one to five matrices; if these matrices are closely homologous, they are grouped in the same cluster (RhizoBindingSites, matrix-clustering section) (Castro-Mondragon et al., 2017). *R. etli* CFN42 contained 15 unique AraC TFs with deduced matrices. Only cluster_99 contained the genes RHE_RS1795 and RHE_RS23180; the other 13 AraC TFs were in different clusters. This indicated that 14 different matrices covered the 15 AraC TFs in this dataset. In contrast, in the S-matrices, the same AraC TFs in cluster_99 were grouped in cluster_65. An additional cluster_12 with the RHE_RS11860, RHE_RS32555, and RHE_RS11860 AraC TFs was shown (Supplementary Table 3A), and the other 12 AraC TFs were grouped into different clusters, suggesting that 14 distinct matrices are involved in the transcriptional regulation of these genes (Supplementary Table 3A). *Rhizobium leguminosarum* bv. *viciae* 3841 with O- and S-matrices had 15 unique AraC TFs. For the O-matrices, cluster_112 contained RL_RS28895 and RL_RS30975 AraCs, whereas cluster_125 contained the RL_RS06995 and RL_RS16545 AraC TFs, which were equally clustered with the S-matrices in cluster_14 and cluster_46, respectively. Eleven other AraC TFs were grouped into different clusters, suggesting that 13 different motifs are involved in the regulation of these genes (Supplementary Table 3B). Moreover, for *S. meliloti* 1021, four unique AraC TFs with both O- and S- matrices were found. For the O matrices, all the AraC TFs were grouped into different clusters. For S-matrices, cluster_3, containing the SMa1454 and SMa2163 AraC TFs were identified (Supplementary Table 3C). Duplication of genes is frequent in species of the Rhizobiales taxon, that is, in *R. etli* CFN42, operons *nifHDK*, *FixNOQP,* and *FixGHIS* (Valderrama et al., 1996) (Granados-Baeza et al., 2007). TFs of the same family may be involved in different metabolic tasks (Cortés-Avalos et al., 2021), which would explain why they are potentially expressed in different metabolic conditions, thus with a different transcriptional regulation, and only some genes with highly homologous matrices may co-occur in their expression (Taboada-Castro et al., 2022). An identical analysis of the matrix-clustering data of the ArsR, GntR, and LysR families with O-matrices was performed to determine whether members of TFs of the same family were clustered in different groups.

#### ArsR family

*R. etli* CFN42, and R. *leguminosarum* bv. *viciae* 3841, and *Sinorhizobium meliloti* 1021 contained eight, five, and five unique ArsR TFs, respectively (Supplementary Table 4A–C). *Rhizobium etli* CFN42 belonged to cluster_172 with RHE_RS05330 and RHE_RS05495 ArsR TFs, and the last six ArsR TFs belonged to different clusters. In contrast, the ArsR TFs were not grouped in the same cluster as R. *leguminosarum* bv. *viciae* 3841 and *Sinorhizobium meliloti* 1021, meaning they have different motif sequences (Supplementary Table 4A–C).

#### GntR family

For the GntR family from *R. etli* CFN42, five clusters were shown (Supplementary Table 5A): cluster_123 with RHE_RS10960 and RHE_RS22955; cluster_3 with RHE_RS23065 and RHE_RS28665; cluster_344 with RHE_RS10960 and RHE_RS24540; cluster_60 with RHE_RS24620 and RHE_RS27280; and cluster_83 with RHE_RS14875, RHE_RS24620, RHE_RS04625, RHE_RS10960, and RHE_RS29975. Ten different GntR TFs were then grouped into these five clusters and the four last GntR TFs from a total of 14 were each grouped in different clusters (Supplementary Table 5A). For R. *leguminosarum* bv. *viciae* 3841, cluster_108 had three genes, and RL_RS35915, RL_RS15070, and RL_RS26800, and cluster_137 with RL_RS15070, and RL_RS29140 were found, totaling four different TFs; then, eight TFs from 12 unique TFs were found in different clusters (Supplementary Table 5B). In *Sinorhizobium meliloti* 1021, cluster_26 contained the SMa0160 and SMa0062 GntR TFs, indicating that seven of the nine unique TFs were located in different clusters (Supplementary Table 5C).

#### LysR family

The most abundant family of TFs was the LysR family in *E. coli* k-12 (Pérez-Rueda and Collado-Vides, 2000) (Perez-Rueda et al., 2018). Matrix-clustering analysis with O-matrices for LysR TFs from *R. etli* CFN42 revealed 15 clusters grouping twenty-seven different genes from a total of forty-two unique LysR TFs with matrices. The remaining 15 LysR TFs were located in different clusters (Supplementary Table 6A). In R. *leguminosarum* bv. *viciae* 3841, twenty-four genes were distributed in 15 clusters, and the last twenty-two of forty-six LysR TFs with matrices were located in different clusters (Supplementary Table 6B). Furthermore, for *Sinorhizobium meliloti* 1021, seven clusters contained 13 LysR TFs, and eleven from a total of twenty-four LysR TFs were grouped into different clusters (Supplementary Table 6C). These data showed that, for O-matrices, TFs from the same family were grouped into different clusters that were potentially subjected to different transcriptional regulations. Consequently, TFs with the same transcriptional regulation should be grouped within the same cluster. For S-matrices in general, a lower number of groups of clusters was found than with O- matrices, meaning that S-matrices were more different than the O-matrices.

### Comparison of O- and S-matrix data with the Regprecise data

To determine the confidence of our data, a comparison of 57 regulons reported in the section on propagated data from Regprecise (Novichkov et al., 2013) and the corresponding h-regulons obtained with the O-matrices in RhizoBindingSites and S-matrices in RhizoBindingSites v2.0 databases were done. In the data on the 57 regulons, the authors did not determine whether the regulons were operons; the numeration of the locus tags and the assignment of the locus names strongly suggest that they were operons. Considering *a priori*, that they are operons, *Rhizobium etli* CFN42 (Supplementary Table 7A), *R. leguminosarum* bv. *viciae* 3841(Supplementary Table 7B), and *Sinorhizobium meliloti* 1021 (Supplementary Table 7C) showed 72.14%, 68.07%, and 81.34% of common genes in the regulons with O-matrices (see the summary in Supplementary Table 7D); meanwhile, 58.96%, 52.10%, and 65.62% of common genes were with S-matrices, respectively. These data show that our predictions coincide with the data from the Regprecise database. Note that these data only have O- and S-matrices that recognize a motif in the upstream regulatory region of TFs, and are expected to be autoregulated TFs. Instead, we noticed that 22.8% of the TFs of regulons from Regprecise were not potentially autoregulated (YrdX, NadQ, NrdR, HutC, Mur, ModE, NifA, HrcA, RhiR, CadR-PbrR, SMc04260, AnsR, and SMb20039); this likely limits the detection of all other genes of regulons from Regprecise. Also, these data showed 13.17%, 15.97%, and 15.72% fewer common genes detected in the S-matrices than the O-matrices for *Rhizobium etli* CFN42 (Supplementary Table 7A), *R. leguminosarum* bv. *viciae* 3841(Supplementary Table 7B), and *Sinorhizobium meliloti* 1021 (Supplementary Table 7C), respectively (see summary in Supplementary Table 7D). Therefore, an additional analysis to determine the number of genes that a TF potentially regulates (TFs hierarchy) in O- and S-matrices for *Rhizobium etli* CFN42, *R. leguminosarum* bv. *viciae* 3841, and *Sinorhizobium meliloti* 1021 showed that, at a p-value of 1.0e-04, 359, 545, and 445, fewer genes were detected on average with S- than O- matrices, respectively. At a p-value of 1.0e-05, on average, 58, 93, and 82 fewer genes were detected in the S- than with the O-matrices, respectively. Meanwhile, at a p-value of 1.0e-06, on average, 6, 9, and 10 fewer genes were detected with S- than with O-matrices, respectively (data not shown). These data agree with the lower detection rate of genes with S-matrices than those with O-matrices of regulons from Regprecise. The difference from the data showing a greater number of unique genes detected with S- than O- matrices (see above) (Supplementary Table 1B) is that, for this analysis, unique genes were considered separately at p-values of 1.0e-04, 1.0e-05, and 1.0e-06, instead of all of them being considered together.

### Conclusions

In the face of global warming, increasing biological constraints are imposed on food production, and the engineering of metabolic pathways to achieve more efficient biological nitrogen fixation in the symbiosis between the Rhizobiales taxon species and their respective host leguminous plants is a desirable strategy. With the availability of genomic sequences, bioinformatics methodologies are essential for extracting pertinent information on transcriptional regulation at the genomic level. Sites, which are conserved short nucleotide sequences located in the upstream regulatory region of genes called motifs, potentially involved in transcriptional regulation, were obtained with the O-matrices deposited in the RhizoBindingSites database. These sites were used to re-deduce new S-matrices. Pointing O-matrices were deduced from the upstream sequences of the orthologous genes of each gene per genome, and S-matrices were deduced from the sites of the genome obtained with the O- matrices. Despite on average, fewer TF genes had S-matrices than O-matrices, a genomic scan analysis with both O- and S- matrices showed, S-matrices had a 1% greater genomic coverage than O-matrices. Globally, these data demonstrate that sequences of S-matrices have more homology with the upstream regulatory sequences of genes than O-matrices in the corresponding genome. Genes in the vicinity detected with S-matrices had a greater TF content than those detected with O-matrices. A hierarchical functional interrelationship was inferred between the transcription factors (Taboada-Castro et al., 2022). Hypothetical regulons were formed with transcription factors grouped using a matrix-clustering method. In addition to this functional validation of the O- and S-matrices, the deduced regulons represented the simplest structure of a transcriptional regulatory network, thus opening the window for the conception of a global transcriptional regulatory network.

This knowledge of the conservation of motifs from symbiotic species, potentially involved in transcriptional regulation, will allow for better designs of experiments to decipher how wiring occurs in a network.

## Author contributions

HT-C, conceptualization, methodology, software, validation, Investigation, data curation, writing original draft, visualization. AJH-A Software, resources, data curation, visualization. JAC-M, conceptualization, methodology, software. SE-G, conceptualization, validation, resources, writing review & editing, supervision, project administration, funding acquisition.

## Acknowledgments

The authors wish to thank the members of the CCG UNAM Information Technology Administration Unit (UATI), for the effort and time they dedicate to the maintenance of the computer system, advice and ho sting of the "RhizoBindingSites” database.

