## Appendix A-D for "RhizoBindingSites v2.0 is a bioinformatic database of DNA motifs potentially involved in transcriptional regulation deduced from sites of the itself genome": Appendix A-D.docx

This is a command, see User´s guide in RhizoBindingSites database

footprint-discovery -v 1 -org Here_nanme_of_the_organism -taxon Rhizobiales -all_genes -sep_genes -lth occ 1 -lth occ_sig 0 -uth rank 50 -return occ,proba,rank -filter -bg_model taxfreq -task query_seq,filter_dyads,orthologs,ortho_seq,purge,dyads,maps,gene_index,index -o name_of_the_output

Appendix B)

This is a Perl program to create matrices from sites:

use strict;

\# module load rsat;

my ($gene_found_flag, $BS_file_list, $folder, $input_file_all_sites, $org) = 0;

my (@BS_file, @BS_file_list) = ();

my (%GeneID_and_BS_file) = ();

#$ENV{"RSAT_BIN"}="/space23/rsat/bin";

\## Organism

$org = "Escherichia_coli_GCF_000005845.2_ASM584v2";

#$org = "Sinorhizobium_meliloti_1021_uid57603";

\## Input file

if($org eq "Escherichia_coli_GCF_000005845.2_ASM584v2"){

$folder = "/home/htaboada/sp27/red_etli/results/matrix-scan/all_genome_E_coli_K12/Rhizobiales/".$org."/1e-3";

print "$folder";

$input_file_all_sites = $folder."/matrix_scan_E_coli_K-12_1e-3.txt";

} elsif($org eq "Sinorhizobium_meliloti_1021_uid57603"){

$folder = "/home/htaboada/red_etli/results/matrix-scan/Rhizobiales_all_matrices/".$org."/1e-3";

$input_file_all_sites = $folder."/matrix_scan_E_coli_K-12_1e-3.txt";

}

my $binding_sites_folder = $folder."/Binding_sites_files";

\## Get the list of files with the sites + Get the gene ID

opendir(my $dir, $binding_sites_folder) or die "Cannot open directory: $!";

my @BS_file_list = readdir $dir;

closedir $dir;

foreach my $file (@BS_file_list){

if($file =~ /^Rhizobiales/){ next;}

if($file =~ /^\.$/){ next;}

if($file =~ /^\.\.$/){ next;}

my @split_line = split(/_all_/, $file);

my $gene_ID = $split_line[0];

$gene_ID =~ s/_purged.fas//;

print "--- $gene_ID --- \n";

$GeneID_and_BS_file{$gene_ID} = $binding_sites_folder."/".$file;

\## Purge sequences

my $purged_seq = $binding_sites_folder."/".$gene_ID."_purged.fas";

my $purge_cmd = "purge-sequence -i ".$GeneID_and_BS_file{$gene_ID}." -format multi -o ".$purged_seq;

print($purge_cmd);

system($purge_cmd);

print "--- $purged_seq--- \n";

\## Dyad-analysis

my $dyad_folder = "/home/htaboada/sp27/red_etli/results/motif_discovery/".$org."/".$gene_ID;

system("mkdir -p ".$dyad_folder);

my $dyad_outfile = "/home/htaboada/sp27/red_etli/results/motif_discovery/".$org."/".$gene_ID."/dyad_analysis_".$gene_ID.".dyads";

my $dyad_command = "dyad-analysis -i ".$purged_seq." -v 2 -quick ";

$dyad_command .= " -sort -type any -2str -noov ";

$dyad_command .= " -lth occ 1 -lth occ_sig 0 -uth rank 50 -l 3 -spacing 0-20 -bg upstream-noorf ";

$dyad_command .= " -org ".$org." -return occ,proba,rank -o ".$dyad_outfile;

system($dyad_command);

\## Pattern-assembly

my $assembly_outfile = "/home/htaboada/sp27/red_etli/results/motif_discovery/".$org."/".$gene_ID."/pattern_assembly_".$gene_ID.".assembly";

my $assembly_command = " pattern-assembly -v 2 -subst 1 -weight 5 -maxfl 1 -toppat 50 -2str -max_asmb_nb 5 -i ".$dyad_outfile." -o ".$assembly_outfile;

system($assembly_command);

\## Convert-matrix

my $matrix_outfolder = "/home/htaboada/sp27/red_etli/results/motif_discovery/".$org."/".$gene_ID."/".$gene_ID."_matrices.tf";

my $matrix_logo_outfolder = "/home/htaboada/sp27/red_etli/results/motif_discovery/".$org."/".$gene_ID."/logos/";

my $matrix_command = " convert-matrix -v 2 -i ".$assembly_outfile." -from assembly -to tf -prefix ".$gene_ID." -return counts,consensus,logo -logo_format png -logo_opt -e -logo_opt -M -logo_dir ".$matrix_logo_outfolder." -o ".$matrix_outfolder;

system($matrix_command);

print "File here: $matrix_outfolder\n";

}

Appendix C)

This is a Perl program to determine vicinity:

open(FINAL,">$ARGV[0].final");

my $deletion = $ARGV[3];

my @POS = split(/:/,$ARGV[2]);

my $ok = 0;

my $c	= 0;

my $d	= 0;

my $index = 0;

my @data = ();

my @num	= ();

my @orden = ();

my @vecinos = ();

open(UNO,$ARGV[0]);

while(<UNO>){

chop;

my $line = $_;

my @A	= split /\t/;

$ok	= 0;

if(/^unicos/){

(@orden) = &ordenar(@num);

for(my $m = 0; $m < @orden; $m++){

#print FINAL $data[$orden[$m]] . "\n";

$vecinos[$m] = $data[$orden[$m]];

}

my $ant = ();

my $sig = ();

my $m	= 0;

my $x	= 0;

my $ext = 0;

my $count = 0;

while($m < @vecinos){

$sig = $vecinos[$m];

if($ant){

($ext) = &extender($ant,$sig,$x);

if($ext == 1){

$vecinos[$m-1] .= "\t$count";

$vecinos[$m] .= "\t$count" if($m == @vecinos - 1);

$ant = $sig;

$x++;

}else{

if($x != 0){

$vecinos[$m-1] .= "\t$count";

$count++;

}

$ant = $sig;

$x = 0;

}

$m++;

}else{

$ant = $sig;

$x = 0;

$m++;

}

}

print FINAL join("\n",@vecinos) . "\n";

$index = 0;

@data = ();

@num	= ();

@orden = ();

}

@vecinos = ();

open(DOS,$ARGV[1]); while(<DOS>){

chop;

my $line2 = $_;

my @B	= split /\t/;

$A[$POS[0]] =~ s/\s+//g;

if($A[$POS[0]] eq $B[$POS[1]]){

$data[$index] = $line2;

$num[$index++] = $B[11];

$c++; $ok=1; last;

}

}close DOS;

if($ok == 0) {

print FINAL "$line\n";

$d++;

}

}close UNO;

close FINAL;

$s = $c + $d;

print "\nFINAL =$c\nRESTO =$d\nSUMA =$s\n";

sub extender{

my ($cA,$cS,$cI) = @_;

my $extend	= 0;

my $type	= 0;

my $replicon = "";

my @A = split(/\t/,$cA);

my @B = split(/\t/,$cS);

$A[5] =~ s/\d+//g;

$B[5] =~ s/\d+//g;

#print "if( (abs($A[9] - $B[9]) <= $deletion) && ($A[5] eq $B[5]) )";<stdin>;

if( (abs($A[11] - $B[11]) <= $deletion) && ($A[5] eq $B[5]) ){

$extend = 1;

}

return ($extend);

}

sub ordenar{

my (@i) = @_;

my @c = ();

my @o = ();

my $ok = 0;

for(my $p = 0; $p < @i; $p++){

$c[$p] = $p;

}

for(my $p = 0; $p < @i; $p++){

if($ok == 0){

$o[0] = $p;

$ok = 1;

}else{

my $pase = 0;

my @a	= ();

for(my $q = 0; $q < @o; $q++){

if($i[$p] >= $i[$o[$q]]){

$a[$q] = $p;

for(my $x = $q; $x < @o; $x++){

$a[$x+1] = $o[$x];

}

$pase = 1;

last;

}else{

$a[$q] = $o[$q];

}

}

if($pase == 0){

$a[@a] = $p;

}

@o = @a;

}

}

return @o;

}

Appendix D)

This is a command to analyze homology between matrices:

matrix-clustering -v 2 -matrix_format transfac -matrix all Here_file_of_matrices_in_transfac_format.tf tf -hclust_method average -calc sum -title Here_a_name -metric_build_tree Ncor -lth w 5 -lth cor 0.6 -lth Ncor 0.4 -label_in_tree name -return json,heatmap -o Here_an_output_file_name -quick &
